# The Conserved CNOT1 Interaction Motif of Tristetraprolin Regulates ARE-mRNA Decay Independently of the p38 MAPK-MK2 Kinase Pathway

**DOI:** 10.1101/2022.02.15.480631

**Authors:** Alberto Carreño, Jens Lykke-Andersen

## Abstract

Regulation of the mRNA decay activator Tristetraprolin (TTP) by the p38 mitogen-activated protein kinase (MAPK) pathway during the mammalian inflammatory response represents a paradigm for the regulation of mRNA turnover by signaling. Phosphorylation of TTP by p38 MAPK-activated kinase 2 (MK2) inhibits the association of TTP with the CCR4-NOT deadenylase complex and represses TTP-mediated mRNA decay. Here we present evidence that TTP remains active in the presence of activated MK2 due to its highly conserved CNOT1 Interacting Motif (CIM), which remains unphosphorylated and capable of promoting deadenylation and decay. The CIM recruits the CCR4-NOT complex cooperatively with previously identified conserved tryptophan residues of TTP and deletion of the CIM strongly represses residual association with the deadenylase complex and activity of TTP in conditions of active MK2. A conserved serine in the CIM is not a target of MK2 but is instead phosphorylated by other kinases including the PKCα pathway and regulates TTP activity independently of MK2. These results suggest that kinase pathways regulate TTP activity in a cooperative manner and that the p38 MAPK-MK2 pathway relies on the activation of additional kinase pathway(s) to fully control TTP function.

## Introduction

mRNA turnover is a critical step in the regulation of gene expression and improper regulation of mRNA stability can promote the development of pathologies including neurodegenerative disorders, cancer, and chronic inflammation (1). mRNA degradation occurs by a multistep process that generally initiates with the removal of the poly(A)-tail, followed by decapping of the 5’ 7-methylguanosine cap, and exonucleolytic decay from either the mRNA 3’ or 5’ end (1, 2). Transcriptome-wide analyses have revealed considerable differences in stability between mRNAs with half-lives in mammals ranging from minutes to hours or days (3, 4). Factors that affect the stability of mRNAs include sequences found within the 3’ or 5’ untranslated regions (UTRs), and the codon usage within the open reading frame (1–3, 5). Importantly, there are transcripts whose stability change in accordance with signaling events in the cell. These transcripts are often regulated by RNA-binding proteins, which are themselves targets of post-translational modification. The general principles by which these post-translational modifications affect the activation of mRNA decay remain poorly defined.

The RNA binding protein Tristetraprolin (TTP; also known as ZFP36 or TIS11) is an mRNA destabilizing factor that recruits factors promoting translation repression, deadenylation, decapping, and exonucleolytic decay (6–9). TTP is a highly regulated protein whose post-translational modifications have been shown to affect its stability, localization, and decay activity (10–13). TTP is best known for its role in resolving the inflammatory response by promoting the degradation of pro-inflammatory cytokine mRNAs that contain adenosine-uridine rich elements (AREs) in their 3’ UTRs (14–18). Structurally, TTP consists of a zinc-finger domain responsible for RNA binding, flanked by amino- (N-) and carboxy- (C-) terminal domains, each of which are capable of promoting mRNA decay (6). Two vertebrate paralogs of TTP, BRF1 (also known as ZFP36L1 and TIS11b) and BRF2 (also known as ZFP36L2 and TIS11d), which also mediate degradation of ARE-containing transcripts, contain zinc finger domains that are highly similar to that of TTP, but N- and C-terminal domains that are distinct, except for a highly conserved region at the extreme C-terminus (10, 19–21).

Recent attention has been turned towards the conserved C-terminal region of the TTP protein family. Studies examining the structural basis of TTP’s association with the central cytoplasmic deadenylase, the CCR4-NOT complex, revealed an interaction with the CCR4-NOT scaffold protein, CNOT1, and this C-terminal region of TTP, which was therefore named the CNOT1-Interacting Motif (CIM) (8, 22). Supporting the importance of the CIM, deletion of the TTP CIM led to a stabilization of target transcripts (8, 23). The loss of the CIM, however, did not completely ablate the ability of TTP to promote mRNA decay, an observation that was further supported by the development of mice lacking the TTP-CIM, which exhibited an inflammatory phenotype significantly milder than that observed upon the complete loss of TTP (23). Separately, it was demonstrated that TTP has a second interaction with the CCR4-NOT complex via several conserved tryptophan residues that interact with the CNOT9 subunit (9). Mutation of those residues were also shown to stabilize TTP target transcripts. Furthermore, TTP is known to interact with the 4EHP-GYF2 translation repression complex and with decapping factors (6, 7, 24). However, the importance of these interactions for TTP’s ability to activate mRNA decay in concert with the CCR4-NOT deadenylase complex remains less well understood.

TTP is a highly post-translationally regulated protein that has been shown to be a target of several kinase pathways. The most well-characterized of these is the p38 MAPK pathway, which is activated during the inflammatory response (25). A downstream kinase in the p38 MAPK pathway, MAPK-activated protein kinase 2 (MK2), targets TTP serine residues 52 and 178 (mouse numbering) whose phosphorylation leads to the stabilization of TTP target transcripts by inhibiting the recruitment of deadenylases (13, 26–28). Furthermore, MK2-mediated phosphorylation of TTP promotes the recruitment of 14-3-3 adaptor proteins, which have been speculated to inhibit degradation factor association, and affect both TTP localization and protein stability (10–12).

Another serine of TTP that undergoes phosphorylation is serine 316 (mouse numbering), a conserved serine of the CIM motif. Phosphorylation of this residue has been shown to inhibit association of TTP-family proteins with CNOT1, suggesting that the association of the major deadenylase complex is repressed by phosphorylation (8, 29, 30). However, a separate study using phosphomimetic mutations, suggested that phosphorylation of the CIM of the TTP homolog BRF1 promotes its association with decapping factors and accelerates mRNA decay (30). While several studies have identified the CIM as a target of MK2 (10, 29, 31), others have implicated the kinases RSK1 and PKA (29, 30). Thus, the importance of the highly conserved CIM and its phosphorylation in TTP regulation and function remains poorly defined.

In this study, to better understand the significance of TTP co-factor interactions in promoting activation of mRNA decay and how these are regulated by phosphorylation, we monitored the combinatorial effects of phosphorylation site and co-factor interaction motif disruptions on TTP function. Our findings show that the CIM acts cooperatively with the conserved tryptophans of TTP to recruit the CCR4-NOT complex and activate mRNA decay, but upon mutation of these motifs, TTP remains unimpaired in its interaction with decapping factors and partially active. Phosphorylation of the TTP CIM reduces its ability to associate with the CCR4-NOT complex and promote mRNA deadenylation, and this phosphorylation event occurs with kinetics similar to that of the MK2-phosphorylated TTP serine 178 in stimulated mouse fibroblasts and macrophage cell lines. However, contrary to what has been previously reported, the CIM is not a target of the MK2 kinase, and TTP remains active in the presence of active MK2, primarily due to the activity of the CIM. Thus, TTP activates mRNA degradation through multiple co-activator complexes and the TTP CIM regulates TTP activity independently of the p38 MAPK-MK2 pathway.

## Results

### The TTP-CIM cooperates with other regions of TTP to promote mRNA decay

To establish a system for monitoring the interplay between the CIM and its phosphorylation with other known functional regions of TTP (Figure 1A), we utilized a previously established MS2-tethering pulse-chase decay assay (32) (Figure 1B) to first test the sufficiency and necessity of the conserved TTP-CIM motif for TTP-mediated mRNA degradation. A tetracycline-inducible β-globin mRNA containing MS2 coat protein binding sites in the 3’UTR was transiently co-expressed in human HeLa Tet-off cells with the MS2 coat protein fused to GFP and the TTP CIM. GFP served as a marker for transfection efficiency, and to stabilize an otherwise unstable MS2 coat protein. The MS2-GFP-CIM fusion protein promoted target mRNA degradation compared to the control MS2-GFP fusion protein alone (Figures 1C and 1D), which was expressed at a similar level (Supplementary Figure S1A). An increase in mobility of the target mRNA through the gel was also observed over time in the presence of the MS2-GFP-CIM fusion protein over the MS2-GFP control (Figure 1C; quantified in Figure 1E), consistent with the previously described function of the CIM in accelerating deadenylation (8). This conclusion was confirmed by oligo-dT and RNase H treatment to remove the poly-A tail, which resulted in the target mRNAs migrating at the same rate (Supplementary Figure S1B). Co-immunoprecipitation (co-IP) assays for the MS2-GFP-CIM fusion protein confirmed CNOT1 association as compared with the control MS2-GFP protein (Supplementary Figure S1C). Thus, our MS2-coat protein tethering system recapitulates previously reported activities of the CIM (8), and the TTP-CIM is sufficient to accelerate mRNA deadenylation and decay.

**Figure 1.**
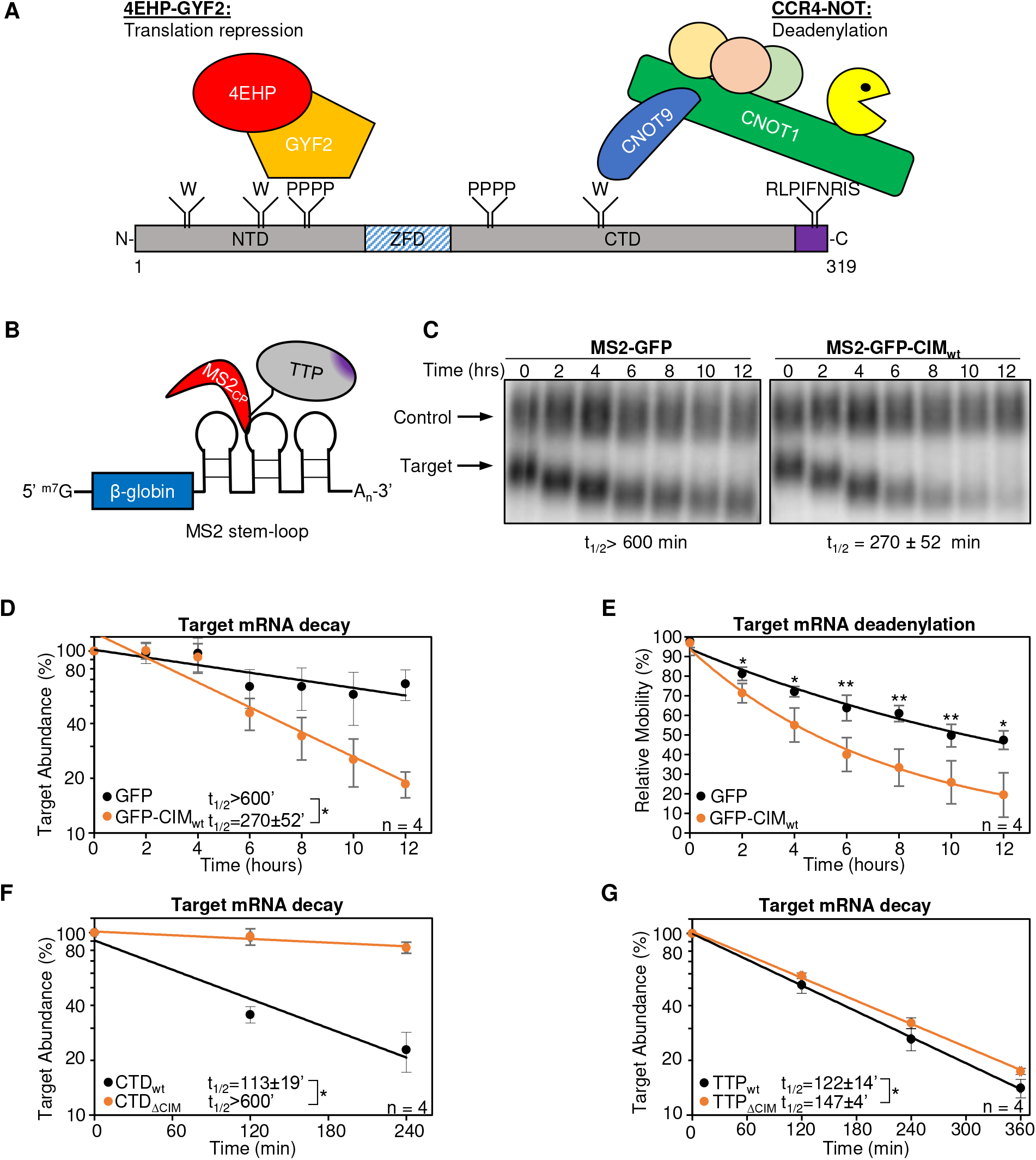
The TTP CIM promotes mRNA decay cooperatively with other regions of the TTP protein. (A) Schematic of mouse TTP, highlighting conserved tryptophan residues (W) interacting with CNOT9, tetraproline motifs (PPPP) interacting with the 4EHP-GYF2 translation repression complex, and the CNOT1-interacting motif (CIM), shown in purple. NTD: N-terminal domain, ZFD: Zinc-Finger Domain, CTD: C-terminal domain. (B) Schematic of the tethered mRNA decay assay. A tetracycline-regulated β-globin mRNA containing MS2 coat protein (MS2cp) stem-loop binding sites in the 3’UTR is targeted by MS2-TTP fusion proteins. (C) Northern blot monitoring the degradation in HeLa Tet-off cells over time after transcriptional shut-off by addition of tetracycline of β-globin mRNA (Target) tethered to MS2-GFP (left) or MS2-GFP-CIM (right) fusion proteins. An extended β-globin mRNA that lacks MS2-coat protein binding sites and whose transcription is not regulated by tetracycline served as an internal control (Control). The half-life of the target mRNA calculated after normalization to the internal control is shown below each panel with standard deviation from four independent experiments. (D) Graph quantifying mRNA decay assays for β-globin mRNA tethered to MS2-GFP fusion proteins shown in panel *C*. Dots represent target mRNA abundance relative to the internal control at each time point with standard deviations shown from four independent replicates (n=4). The curves represent best fits to first-order degradation. Calculated half-lives are given with standard deviation. *: p<0.05; student’s two-tailed t-test. (E) Graph showing relative band mobilities as a measure of mRNA deadenylation from the experiments in panel *C*. Dots represent mobility of the target mRNA relative to the control mRNA, with the mobility at time 0 set to 100% and the mobility of a deadenylated target mRNA, generated by treatment with oligo-dT and RNase H (Supplementary Figure S1B), set as 0%. Error bars represent standard deviation. *: p<0.05, **: p<0.01 (two-tailed student’s t-test, comparing band mobility at each time point). (F) Target β-globin mRNA degradation, graphed as in panel *D*, in the presence of tethered MS2-CTD_wt_ (black) or MS2-CTD_ΔCIM_ (orange). (G) Same as panel *F*, monitoring degradation of target mRNA tethered to MS2-TTP_wt_ (black) or MS2-TTP_ΔCIM_ (orange).

Although the TTP-CIM can promote mRNA decay on its own, other regions of TTP have been implicated in mRNA decay as well. We were therefore interested in understanding the importance of the CIM in promoting mRNA degradation in conjunction with other functional motifs of TTP. We first deleted the CIM from full-length TTP and from the carboxy-terminal domain (CTD) of TTP, each fused to the MS2 coat protein, and tested the effect on mRNA degradation. Consistent with previous reports (6), the target mRNA was rapidly degraded upon tethering of both the TTP-CTD and full-length TTP (Figures 1F and 1G and Supplementary Figures 2A-D). Deletion of the CIM greatly stabilized target transcripts in the context of the CTD (Figure 1F), while it only marginally impaired the activity of full-length TTP (Figure 1G). This difference in the contribution of the CIM to mRNA decay promoted by the TTP-CTD and full-length TTP suggests a redundancy of the CIM with other regions of TTP, particularly outside of the CTD.

### The TTP-CIM cooperates with conserved tryptophan residues to promote mRNA deadenylation and decay

TTP was previously reported to interact with another component of the CCR4-NOT complex, CNOT9, through its conserved tryptophan residues (Figure 1A) (9). Therefore, we were interested in understanding to what extent the CIM and tryptophan residues cooperate to activate mRNA decay. We generated alanine mutants for all conserved tryptophan residues of TTP and the TTP-CTD with the additional mutation of Proline 257, which was previously shown to also contribute to CNOT9 association (9) (Supplementary Table 1). We performed tethered pulse-chase mRNA decay assays for these mutants with and without CIM deletion. The tryptophan to alanine (WA) mutation caused no significant reduction in activity in the context of the TTP-CTD with or without CIM deletion (Figure 2A and Supplementary Figures S2A and S2B). However, for full-length TTP, further stabilization of the target mRNA was observed when the removal of the CIM was combined with mutations in the tryptophan motifs (Figure 2B and Supplementary Figures S2C and S2D). This reduction in activity was accompanied by reduced association of TTP and the TTP CTD with CNOT1 and CNOT9 components of the CCR4-NOT complex upon mutation of CIM and tryptophan motifs (Figures 2C and D). These observations demonstrate that the conserved tryptophan motifs and the CIM serve to cooperatively recruit the CCR4-NOT complex and activate mRNA decay, likely via their previously reported interactions with different subunits of the CCR4-NOT complex.

**Figure 2.**
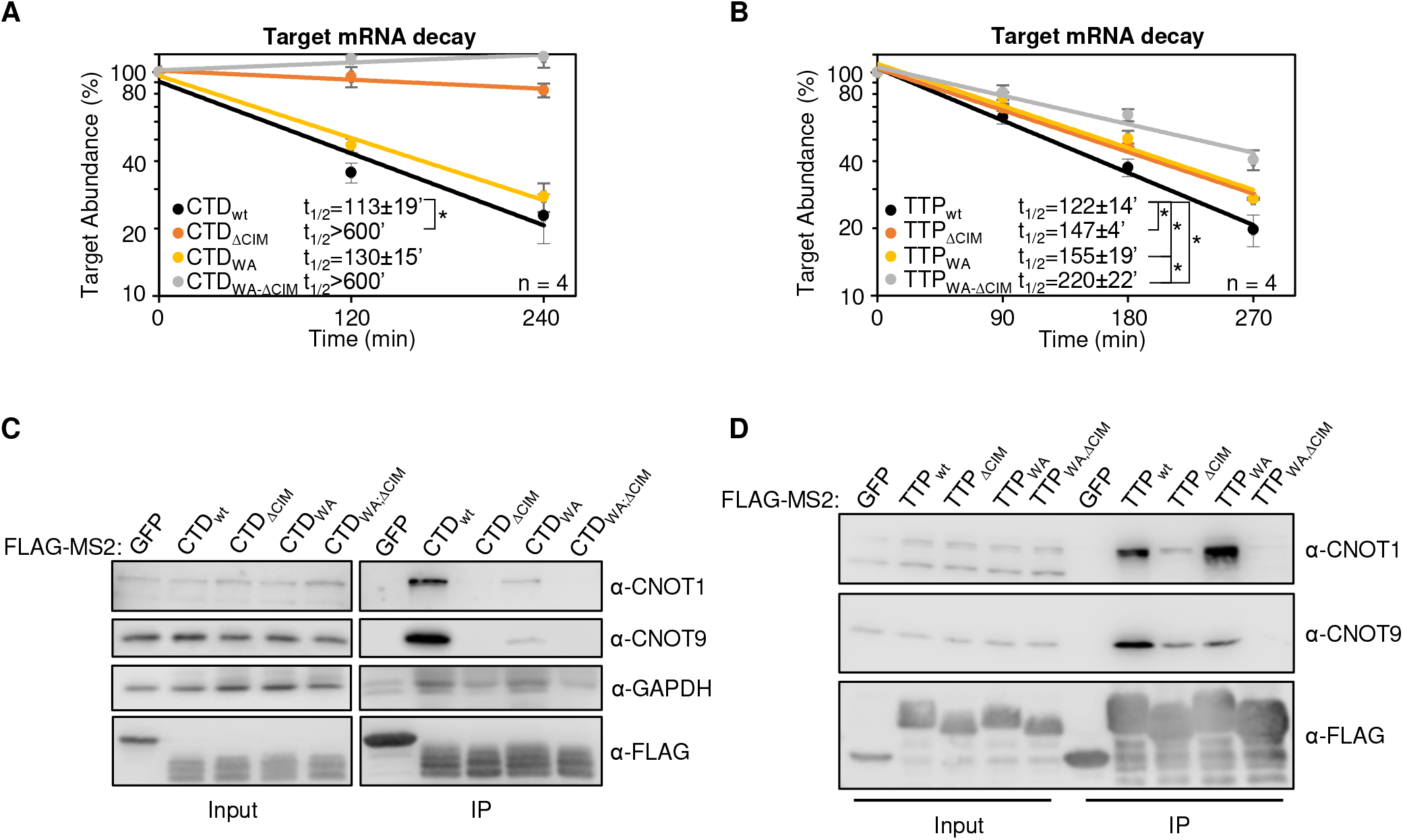
Cooperative activation of deadenylation by the TTP CIM and conserved tryptophans. (A) mRNA decay assays for β-globin mRNA tethered to TTP-CTD wild-type (wt) or mutant proteins with the CIM deleted (ΔCIM), conserved tryptophans mutated to alanines (WA), or both (WA,ΔCIM), graphed as in Figure 1D. *: p<0.05; student’s two-tailed t-test. (B) Same as panel *A*, but with wild-type or mutant MS2-TTP full-length fusion proteins. (C) Western blots showing proteins co-immunoprecipitating (IP, right panels) with the indicated FLAG-tagged MS2-CTD fusion proteins from HEK293T cells after treatment with RNase A, as compared with input samples (left). IP samples correspond to 2.5% of the input. (D) Same as panel *C*, but monitoring proteins associated with FLAG-tagged MS2-TTP fusion proteins.

### Mutation of conserved tetraproline motifs does not significantly affect TTP-mediated mRNA decay rates

Other regions of TTP that interact with mRNA repression factors are the tetraproline motifs, which are conserved among TTP orthologs and promote the association with the 4EHP-GYF2 translation repression complex (7, 33). Mutations in these motifs, and depletion of 4EHP, were previously found to cause increased protein production from TTP target transcripts. Given the interconnected roles of translation repression and mRNA decay, we next performed tethered mRNA decay assays for TTP tetraproline mutants. We found that mutation of TTP tetraproline motifs (PS) did not increase the stability of the tethered target mRNA (Figure 3A and Supplementary Figure S3A). To address whether the tetraproline motifs act cooperatively with the CNOT1 and CNOT9 interaction motifs, we established double and triple mutant TTP proteins. Comparing TTP mutants with and without tetraproline motif mutations revealed no stabilization of the target mRNA attributed to the tetraproline mutations (PS) when combined with the tryptophan mutations (WA) and/or CIM deletion (ΔCIM) (Figures 3B and 3C and Supplementary Figures S3A-D), despite loss of association with CCR4-NOT and GYF2-4EHP complexes as monitored by co-immunoprecitation assays for CNOT1, CNOT9 and GYF2 (Figure 3D). Interestingly, the TTP triple mutant protein defective in both CCR4-NOT and GYF2-4EHP complex association maintained undisrupted association with DDX6 and EDC4 of the decapping complex (Figure 3D), suggesting that additional functional motifs exist in TTP that may explain the partial activity of this triple mutant.

**Figure 3.**
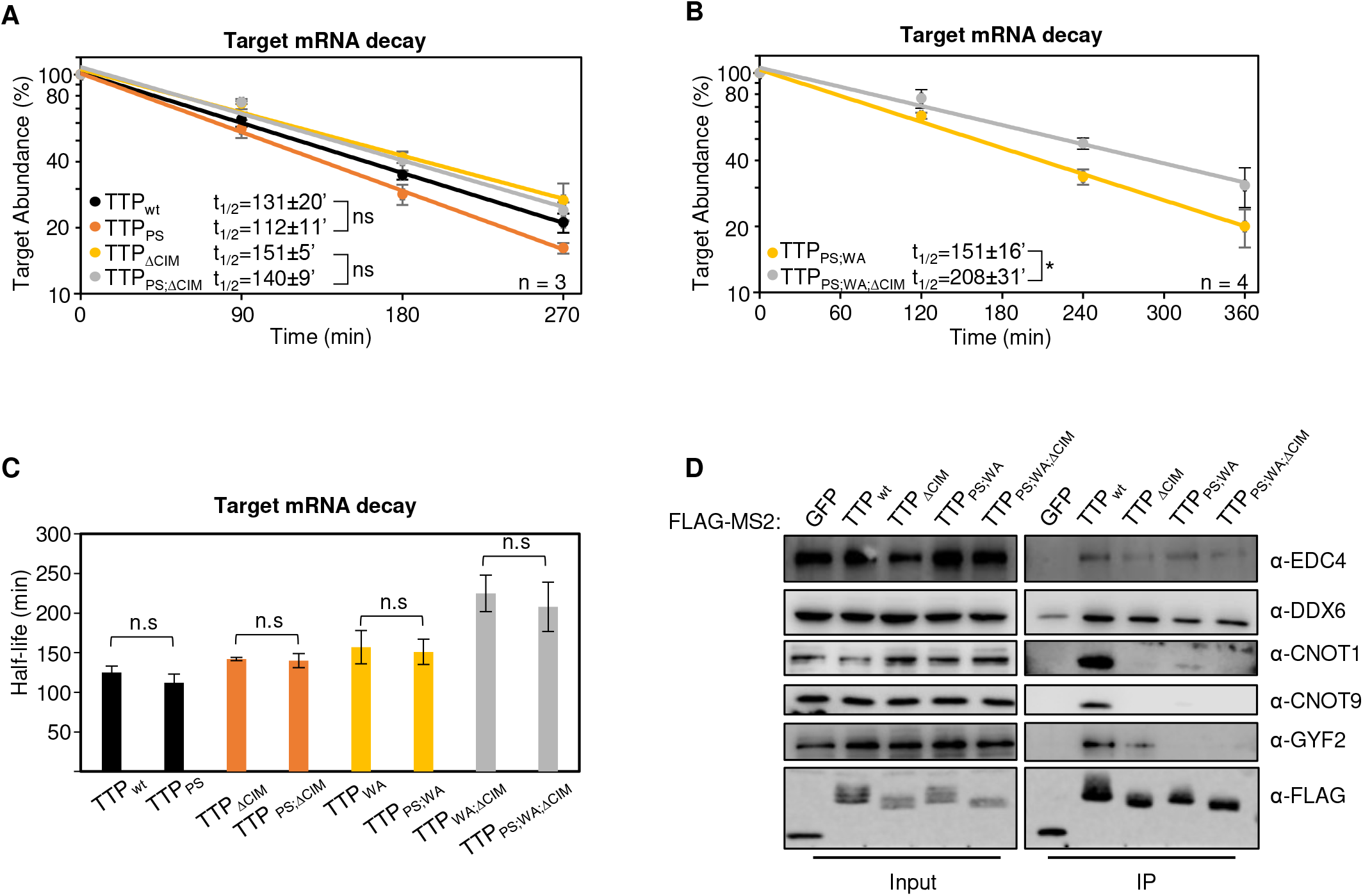
TTP tetraproline motifs are not rate-limiting for mRNA decay. (A) mRNA decay assays for β-globin mRNA tethered to indicated MS2-TTP wild-type (wt) or mutant proteins with the tetraproline motifs mutated to serines (PS), the CIM deleted (ΔCIM), or both (PS,ΔCIM), graphed as in Figure 1D. n.s: p>0.05 (student’s two-tailed t-test). (B) Same as panel *A*, with indicated mutant MS2-TTP fusion proteins with tetraproline motifs mutated to serines and conserved tryptophans to alanines (PS,WA), or additional deletion of the CIM (PS,WA,ΔCIM). *: p<0.05; student’s two-tailed t-test. (C) Bar-graph comparing half-lives of β-globin mRNA tethered to the indicated MS2-TTP fusion proteins with or without mutation in tetraproline motifs (PS). n.s: p>0.05 (student’s two-tailed t-test). (D) Western blots showing proteins co-immunoprecipitating (IP, right panels) with indicated FLAG-tagged MS2-TTP fusion proteins from HEK293T cells after treatment with RNase A, as compared with input samples (left). IP samples correspond to 2.5% of the input.

### TTP is transiently phosphorylated at the conserved CIM serine residue

Having established the degree of cooperativity between the TTP CIM and other known functional motifs of TTP, we next turned to the significance of CIM phosphorylation. We generated and validated a phospho-specific antibody raised against the phosphorylated serine 316 residue of mouse TTP (Supplemental Figure S4). Consistent with a recent report (29), induction of the inflammatory response by serum shock of mouse NIH3T3 fibroblasts (Figure 4A), or treatment with lipopolysaccharide (LPS) of mouse RAW264.7 macrophages (Figures 4B), both caused rapid upregulation of TTP accompanied by phosphorylation at TTP serine 316, with kinetics similar to what has been described previously for TTP serine 178 phosphorylation (10, 28).

**Figure 4.**
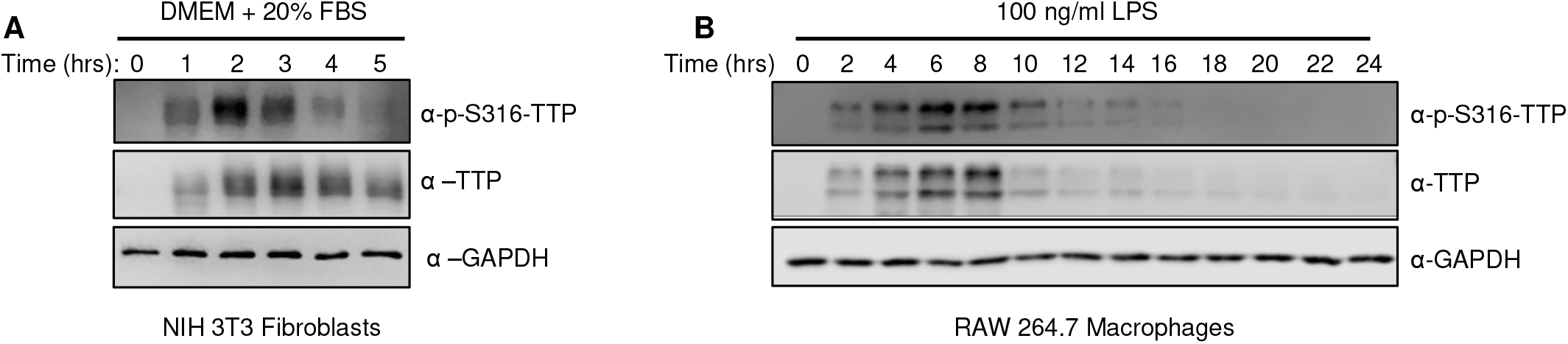
TTP serine 316 is phosphorylated during TTP induction. (A) Western blots monitoring levels of TTP and its phosphorylation at serine 316 (p-S316-TTP) in serum-shocked mouse NIH 3T3 cells. GAPDH serves as a loading control. (B) Same as panel *A*, but monitoring TTP in LPS-induced RAW 264.7 mouse macrophage cells.

### The TTP-CIM is phosphorylated in a p38 MAPK-MK2-independent manner

The TTP-CIM has been reported as a target of several kinase pathways (10, 29–31), including the p38 MAPK-MK2 pathway, which also targets TTP serines 52 and 178 (10, 28). HeLa cells transiently expressing TTP show detectable levels of phosphorylation of both serines 178 and 316 of TTP (Figure 5A). Treatment with the p38 MAPK inhibitor SB203580 (34) decreased serine 178 phosphorylation, as expected, while surprisingly serine 316 phosphorylation remained unaffected (Figure 5A). Consistent with this observation, co-expression of TTP or the TTP-CIM in HEK293 cells with constitutive active MK2 (12, 28) resulted in phosphorylation of TTP serine 178 (Figure 5B) but not serine 316 (Figures 5B and 5C).

**Figure 5.**
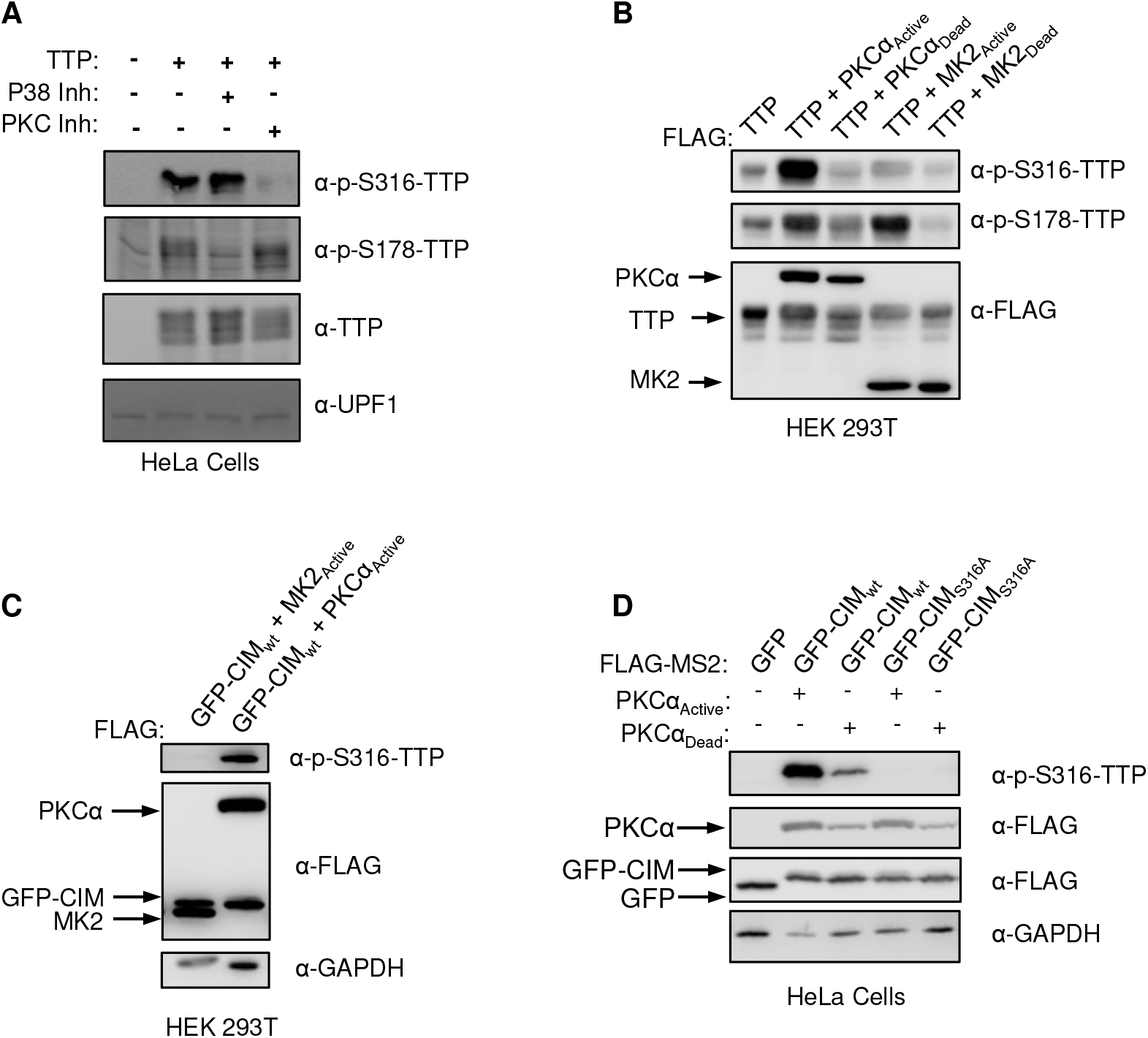
TTP serine 316 is phosphorylated by kinase(s) other than MK2. (A) Western blots monitoring TTP and its phosphorylation at serine 316 (p-S316-TTP) or serine 178 (p-S178-TTP), exogenously expressed in HeLa cells incubated with inhibitors of p38 (SB203580) or PKC (Gö 6983) kinases. The left lane is a control sample from HeLa cells not expressing TTP. UPF1 served as a loading control. (B) Western blots monitoring phosphorylation of exogenous FLAG-tagged TTP co-expressed with FLAG-tagged, constitutive active or catalytic dead, MK2 or PKCα kinases in HEK293T cells. (C) Western blots monitoring phosphorylation of FLAG-tagged GFP-CIM fusion protein co-expressed with indicated constitutive active kinases in HEK293T cells. (D) Western blots monitoring phosphorylation of FLAG-tagged GFP-CIM and GFP-CIM S316A mutant fusion proteins co-expressed with constitutive active or catalytic dead PKCα kinase in HeLa cells.

Other kinases that have been observed to phosphorylate the CIM of TTP-family proteins include p90 ribosomal S6 kinase 1 (RSK1) and Protein Kinase A (PKA) (29, 30, 35). Another kinase predicted to phosphorylate TTP serine 316 in a kinase prediction algorithm (36) is Protein Kinase C-alpha (PKCα). Treatment of HeLa cells with a PKC inhibitor, Gö6983 (37), decreased TTP Serine 316 phosphorylation while Serine 178 phosphorylation remained unaffected (Figure 5A). Moreover, constitutive active PKCα increased Serine 316 phosphorylation both in the context of full-length TTP (Figure 5B) and the TTP-CIM (Figures 5C and 5D). We also observed an increase in TTP serine 178 phosphorylation in the presence of active PKCα (Figure 5B), which is likely due to reported activation by PKCα of the p38 MAPK pathway (38). Thus, serine 316 of the CIM can be phosphorylated by kinases including PKCα, but unlike serines 52 and 178, is not a target of MK2.

### MK2-phosphorylated TTP promotes mRNA decay via the CIM

TTP activity has been previously reported to be inhibited by MK2 phosphorylation on serine residues 52 and 178, correlating with decreased association with deadenylation factors as well as the recruitment of 14-3-3 adaptor proteins (10–12, 39). Our observation that the CIM is not a target of MK2 phosphorylation raised the possibility that the CIM continues to promote deadenylation and decay in the presence of active MK2. To test whether the CIM indeed acts independently of MK2 phosphorylation, we performed tethered mRNA decay assays comparing the effect of MK2 on wild-type TTP and TTP deleted of the CIM. Consistent with previous reports (12, 28), constitutive active MK2 (MK2A) stabilized the TTP target transcript, as compared to catalytic dead MK2 (MK2D), although this effect was relatively small (Figure 6A and Supplementary Figures S5A and S5B). However, removal of the CIM from TTP, while having only a minor effect on TTP activity in the presence of inactive MK2, dramatically stabilized the target mRNA in the presence of active MK2 (Figure 6A and Supplementary Figures S5A and S5B). This stabilization was dependent on the two serines, serines 52 and 178, targeted by MK2 as mutating them to alanines (TTP-2A) restored the activity of TTP deleted of the CIM in the presence of active MK2 (Figure 6B and Supplementary Figures S5C and S5D). Consistent with these observations, treatment of TTP with the phosphatase inhibitor okadaic acid, which causes general upregulation of TTP phosphorylation, caused a reduction in association of TTP with CNOT1, which was exacerbated when the CIM was deleted (Figure 6C). We observed similar strong dependence on the CIM of TTP activity in the presence of active MK2 when the activity of the TTP_PS;WA;ΔCIM_ triple mutant was compared to TTP_PS;WA_ and when wild-type TTP was compared to TTP mutated at a phenylalanine residue within the CIM critical for association with CNOT1 (Supplementary Figures S5E-G).

**Figure 6.**
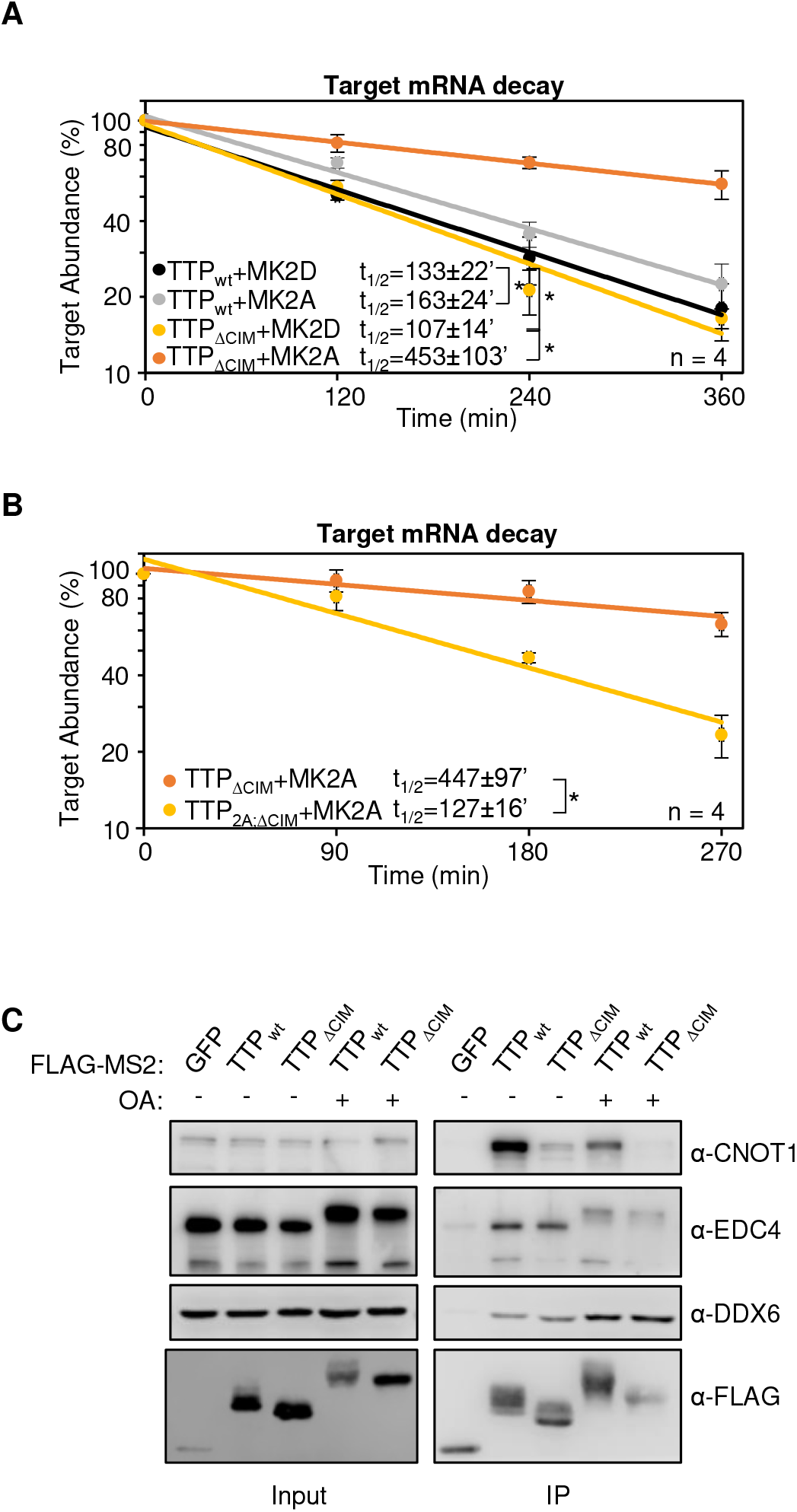
The TTP-CIM remains active in the presence of active MK2. (A) mRNA decay assays for β-globin mRNA tethered to MS2-TTP wild-type (wt) or ΔCIM proteins, co-expressed with constitutive active (MK2A) or catalytic dead (MK2D) MK2 kinases and graphed as in Figure 1D. *: p<0.05; student’s two-tailed t-test. (B) mRNA decay assays for β-globin mRNA tethered to MS2-TTP ΔCIM protein or MS2-TTP ΔCIM with alanine mutations at serines 52 and 178 targeted by MK2 (2A, ΔCIM), co-expressed with constitutive active MK2 (MK2A). (C) Western blot showing proteins co-immunoprecipitating (IP, right panels) with indicated FLAG-tagged TTP proteins expressed in HEK293T cells treated with or without okadaic acid. Input samples corresponding to 2.5% of IPs are shown on the left.

### The TTP-CIM is regulated by phosphorylation independently of MK2

Phosphorylation of the TTP CIM has been previously observed to inhibit association with CNOT1 (8), while another report focusing on BRF1 observed increased association with a decapping factor (30). We therefore wished to test whether CIM phosphorylation regulates TTP activity independently of MK2. To first test the importance of the phosphorylated serine 316 residue in mRNA decay activated by the TTP-CIM alone, we mutated this residue to an alanine, which prevented CIM phosphorylation (Supplementary Figure S1A). MS2-tethered pulse-chase decay assays showed an increase in the degradation rate as a result of the serine to alanine mutation (Figure 7A and Supplementary Figure S6A), and an apparent corresponding increase in deadenylation (Figure 7B), although the latter did not reach a level of statistical significance. Expression of PKCα caused an acceleration of mRNA degradation that was independent of TTP-CIM phosphorylation (Supplemental Figure S7B) and could therefore not be used as a means to test the effect of TTP-CIM phosphorylation. We next tested the importance of CIM phosphorylation in the context of full-length TTP. Mutating serine 316 to an alanine, which prevented phosphorylation of the CIM (Figure 7C), resulted in a minor increase in the rate of degradation by TTP in the presence of inactive MK2. This effect of the serine 316 to an alanine mutation was further exacerbated in the presence of active MK2 (Figure 7D) consistent with TTP relying on an unphosphorylated CIM to activate mRNA decay in the presence of active MK2. Altogether, these findings demonstrate that the TTP CIM activates mRNA decay unregulated by the p38 MAPK-MK2 pathway and that the two separate events of MK2-mediated phosphorylation and phosphorylation of the CIM acts cooperatively to regulate TTP activity.

**Figure 7.**
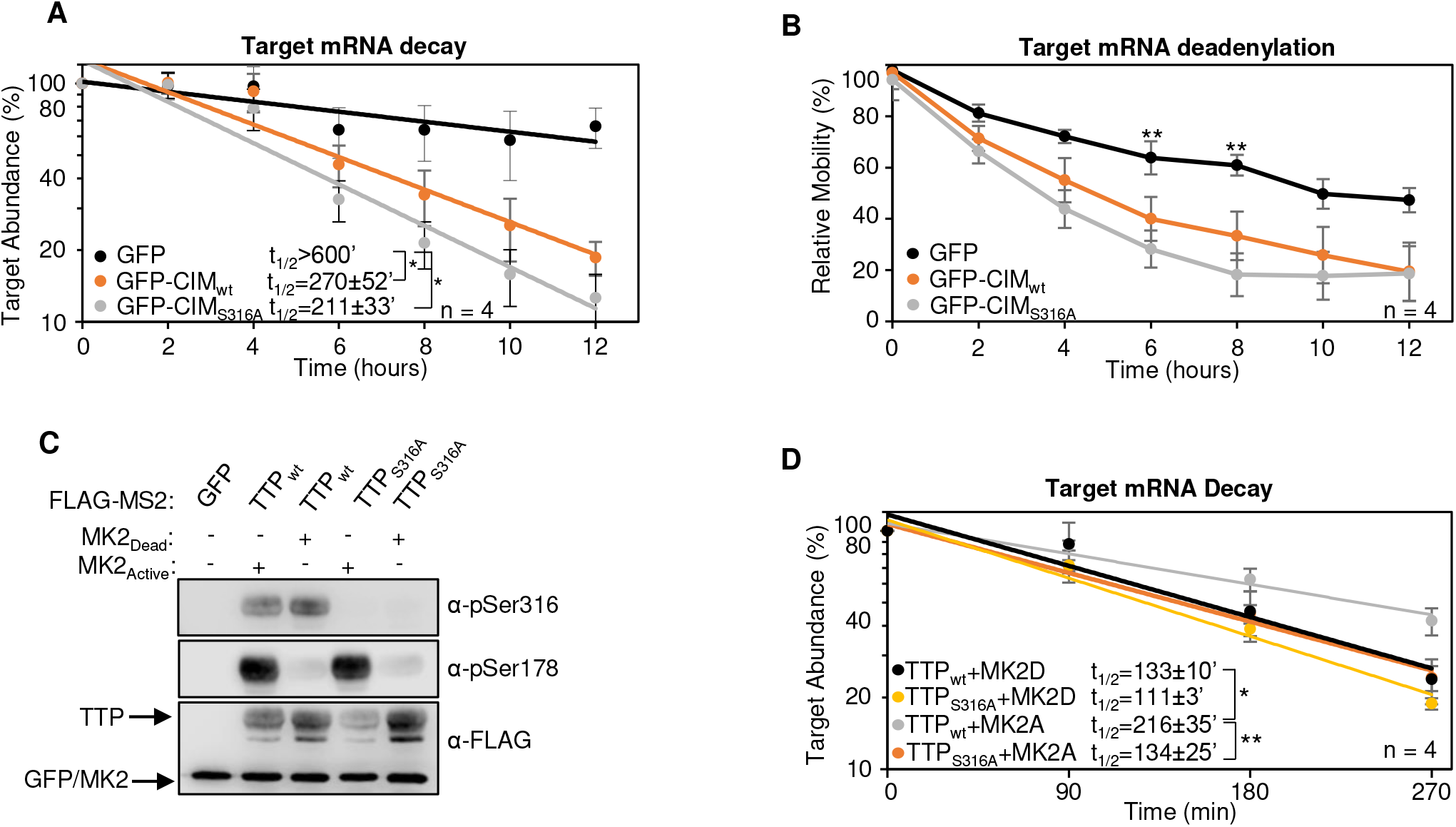
TTP-CIM activity is regulated by phosphorylation independently of MK2. (A) mRNA decay assays for β-globin mRNA tethered to indicated MS2-GFP fusion proteins in HeLa Tet-off cells. *: p<0.05; student’s two-tailed t-test. (B) Graph showing relative band mobility as a measure of mRNA deadenylation from the experiments in panel *A*, measured as described in Figure 1E (see also Supplementary Figure S6). **: p<0.01; student’s two-tailed t-test. (C) Western blots monitoring phosphorylation of MS2-TTP wild-type (wt) or S316A mutant fusion proteins in the presence of constitutive active (MK2A) or catalytic dead (MK2D) MK2 kinase in HeLa Tet-off cells. (D) mRNA decay assays for β-globin mRNA tethered to MS2-TTP proteins co-expressed with MK2 kinases described in panel *C*. *: p<0.05, **: p<0.01; student’s two-tailed t-test.

## Discussion

The regulation of TTP by the p38 MAPK-MK2 pathway during the inflammatory response is a well-established paradigm for the regulation of mRNA decay by signaling. In this study, we demonstrate that the highly conserved C-terminal CIM motif of TTP, which plays a key role in connecting TTP family proteins with the CCR4-NOT deadenylase complex central to mRNA decay, is regulated independently of the p38 MAPK-MK2 pathway. This conclusion is supported by our observations that, unlike TTP serine 178, the TTP CIM serine 316 remained phosphorylated in the presence of a p38 MAPK inhibitor and was not phosphorylated by constitutive active MK2 (Figure 5). Moreover, TTP remained partially active in the presence of constitutive active MK2, largely due to the activity of the CIM (Figure 6). The CIM, in turn, is regulated by other kinases, such as PKCα (Figure 5), RSK1 (29, 35), and PKA (30), whose phosphorylation of the CIM results in impaired association with CNOT1 of the CCR4-NOT complex and reduced decay activity (Figure 7) (8, 31). The TTP CIM recruits the CCR4-NOT complex, and activates deadenylation and decay, in cooperation with conserved tryptophans of TTP (Figures 1 and 2), which had previously been found to interact with CNOT9 (9). By contrast, association of TTP with the 4EHP-GYF2 complex via TTP tetraproline motifs was not rate-limiting for TTP-mediated mRNA decay (Figure 3).

Our finding that TTP remains highly active even when deletion of the TTP CIM and mutations in conserved tryptophans and tetraproline motifs disrupt associations with CCR4-NOT and 4EHP-GYF2 complexes (Figures 1-3), suggests that TTP promotes degradation via additional interactions with cellular RNA decay machinery. This residual activity of TTP may in part be explained by its undisrupted association with decapping factors (Figure 3D). With the exception of its conserved RNA binding zinc-finger domain, TTP is composed of mostly predicted intrinsically disordered regions (23). These types of domains have been implicated in the association with decapping factors and p-bodies and therefore may contribute to the activity of TTP (40–42). It is also possible that additional regions of TTP contribute to the association with the CCR4-NOT complex or other, yet to be determined, mRNA decay factors. Our mutational studies found no rate-limiting role for the 4EHP-GYF2 complex in mRNA decay by tethered TTP. 4EHP-GYF2 may specifically promote translation repression (7, 24, 33). The knockout of 4EHP in mouse embryonic fibroblasts caused upregulation of TTP target mRNAs in addition to their encoded proteins (Fu et al. 2016), which could be an indirect effect of interfering with TTP-mediated translation repression during inflammation. An important goal for future studies will be to identify the complete interaction network of TTP with mRNA decay and translation repression machinery.

The most surprising finding of this study was that the TTP CIM remains active and capable of promoting mRNA decay in the presence of active MK2, despite the p38 MAPK-MK2 pathway having been previously implicated in TTP CIM phosphorylation and inhibition of TTP deadenylase association (28, 29, 31, 34, 43, 44). This observation suggests that activation of the p38 MAPK-MK2 pathway is insufficient to fully inactivate TTP and cooperativity with one or more additional kinase pathways is required. This has important implications for how TTP activity is repressed in physiological conditions, for example during the inflammatory response when TTP repression allows accumulation of ARE-mRNAs expressing pro-inflammatory cytokines (14–16, 18). The observation that TTP associates with the CCR4-NOT complex via additional interactions beyond the CIM (9) (Figure 2) may explain how phosphorylation by MK2 of residues outside of the CIM impairs CCR4-NOT association (26, 28–30). The CIM and its phosphorylated serine is a highly conserved motif shared by all members of the ZFP36 family. ZFP36 paralogs have both overlapping as well as unique roles in mRNA decay. Thus, certain kinase pathways may regulate all members of the ZFP36 family, for example via phosphorylation of the conserved CIM, whereas other kinase pathways may specifically regulate a subset of ZFP36 family members. This may provide specificity to how ARE-mRNA decay is regulated in different cell types and under different conditions.

## Materials and Methods

### Plasmids

pcDNA3-FLAG-MS2 plasmids expressing FLAG-MS2-tagged TTP mutant proteins (Supplementary Table 1) were generated from previously described pcDNA3-FLAG-MS2-TTP plasmids (28) using site-directed mutagenesis according to manufacturer’s protocol (New England Biolabs). Expression plasmids for FLAG-tagged constitutive active and catalytically dead MK2 kinases were previously described (12, 28). Similarly, expression plasmids for the control β-globin mRNA containing a 3’UTR sequence from GAPDH mRNA and the target tetracycline-regulated β-globin mRNA containing a 3’UTR with six MS2 coat protein stem-loop binding sites were previously described (32). The pcDNA3-FLAG-MS2-GFP-CIM constructs were generated by first generating a pcDNA3-FLAG-MS2-CIM plasmid by inserting the sequence corresponding to mouse TTP residues 304-319 into the pcDNA3-FLAG-MS2 plasmid (6) between restriction sites *Bam*HI and *Not*I. Gibson Assembly (New England Biolabs (NEB)) was subsequently performed to insert the GFP coding region from pSpCas9(BB)-2A-GFP (PX458) into the pcDNA3-FLAG-MS2-CIM vector between the MS2 coat protein and the CIM. pcDNA3-FLAG-PKCα constructs expressing FLAG-tagged PKCα were generated by amplifying the coding region of PKCα from HeLa Tet-off cell total RNA and inserting it into pcDNA3-FLAG (45) immediately downstream of the FLAG peptide sequence using Gibson Assembly (NEB). Constitutive active PKCα-AE was generated by mutating alanine 25 into glutamic acid. Catalytic dead PKCα-KR was generated by mutating lysine 368 into arginine.

### Cell Culture

HeLa Tet-off, HEK293T, RAW264.7, and NIH3T3 cells were cultured in Dulbecco’s modified Eagle’s medium (DMEM; Gibco) with 10% fetal bovine serum (FBS) and 1% Penicillin-Streptomycin. For experiments shown in Figure 4A, NIH3T3 cells were washed with phosphate-buffered saline (PBS) and grown in DMEM containing 0.5% FBS for 24 hours. Cells were subsequently treated with DMEM containing 20% FBS for the indicated amount of time and collected in Laemmli sample lysis buffer (2% SDS, 10% Glycerol, 60 mM Tris-HCl pH 6.8, 5% β-Mercaptoethanol). For experiments shown in Figure 4B, RAW264.7 cells were treated with DMEM containing 100 ng/mL lipopolysaccharide (LPS) for the indicated times and collected in sample lysis buffer.

### Antibodies

Rabbit polyclonal anti-pS316-TTP antibody was generated with the synthesized oligo peptide PRRLPIFNRI(p)SVSE (Pocono Rabbit Farm & Laboratory, USA). The antiserum was purified by affinity chromatography by first collecting antibody binding to a PRRLPIFNRI(p)SVSE-peptide column, and, after elution, collecting the flow-through from a subsequent unphosphorylated PRRLPIFNRISVSE-peptide column. Western blots were probed with the following antibodies at the indicated concentrations: rabbit polyclonal anti-FLAG (Millipore Sigma, F7425; 1:1,000), mouse monoclonal anti-FLAG M2 (Millipore Sigma, F1804; 1:1000), rabbit polyclonal anti-DDX6 (Bethyl, A300-461A; 1:1,000), rabbit polyclonal anti-GIGYF2 (Santa Cruz Biotechnology, sc-134708; 1:50), mouse monoclonal anti-14-3-3 (Santa Cruz Biotechnology, sc-1657; 1:1,000), rabbit polyclonal anti-TTP (Sigma-Aldrich, T5327; 1:500), rabbit polyclonal anti-CNOT1 (Proteintech, 14276-1-AP; 1:200), rabbit polyclonal anti-CNOT9 (Proteintech, 22503-1-AP, 1:500), mouse monoclonal anti-GAPDH (Cell Signaling Technology, 97166, 1:1,000), rabbit polyclonal anti-EDC4 ((42); 1:200) rabbit polyclonal anti-pSer178-TTP ((11) a generous gift from Dr. Georg Stoecklin, Heidelberg University; 1:500).

### Co-Immunoprecipitation assays

Co-immunoprecipitation assays were performed as previously described (7). Briefly, confluent HEK 293T cells were split at 1:10 onto 10-cm plates. The following day, samples were transfected with 5 μg of the indicated FLAG-tagged TTP wild-type or mutant expression plasmids using TransIT 293 reagent according to the manufacturer’s protocol (Mirus). 48 hours after transfection, samples were washed in PBS, pelleted by centrifugation, and collected in hypotonic lysis buffer (10 mM Tris-HCl pH 7.5, 10 mM NaCl, 2 mM EDTA, 0.5% triton X-100, 1 mM PMSF, 1 μM aprotinin, and 1 μM leupeptin) for 10 minutes on ice. NaCl concentrations were increased to 150 mM and samples were treated with 50 μg/mL RNase A for 10 minutes. Samples were centrifuged at 21,130*g* for 15 minutes at 4°C and supernatants were added to 50 μl of M2 anti-FLAG-agarose beads (Sigma). Beads were washed 4 times with NET-2 (50 mM Tris-HCl pH 7.5, 150 mM NaCl, 0.05% Triton-X100) and resuspended in 50 μl of 2x Laemmli sample lysis buffer (4% SDS, 20% Glycerol, 120 mM Tris-HCl pH 6.8, 10% β-Mercaptoethanol). Samples were separated by SDS-polyacrylamide gel electrophoresis (SDS-PAGE) and analyzed by western blot. 0.5% of whole cell extracts and 20% of pull-down elutions were loaded for analyses.

### Pulse-chase mRNA decay assays

Tethered mRNA decay assays were performed as previously reported (28, 32). Briefly, confluent HeLa Tet-off cells (Takarabio) in 10-cm plates were split into 12-well plates at between 1:15 to 1:20. Two days later, cells were transfected with β-globin control (25 ng) and reporter constructs containing MS2 binding sites (250 ng) and FLAG-MS2-TTP constructs (100 ng), or MS2-GFP constructs as a control, using TransIT-HeLaMONSTER transfection kit according to the manufacturer’s protocol (Mirus). Cells were subsequently incubated in the presence of 50 ng/ml tetracycline to inhibit expression of the target mRNA. 24 hours post-transfection, transcription of the reporter transcripts was pulsed with the addition of fresh media lacking tetracycline for 4-6 hours and then treated with 1 μg/ml tetracycline to shut off transcription. Total RNA was harvested at indicated times after transcription shut-off by extraction with Trizol following the manufacturer’s protocol (Thermo Fisher) and analyzed by Northern blotting as previously described (28). Deadenylated samples were generated by resuspending total RNA in 9 µl water and incubating the samples with 1 μl of 1 mM oligo-dT_20_ at 80°C for 2 minutes followed by annealing at room temperature for 5 minutes. 7 μl of water, 2 μl of 10x RNAse H Buffer (500 mM Tris-HCl, 750mM KCl, 30mM MgCl2, 100mM DTT, pH 8.3; NEB) and 1 μl of 5 units/µl RNase H (NEB) were added and samples were incubated at 37°C for 30 minutes. RNA samples were extracted with phenol:chloroform (1:1), followed by ethanol precipitation, and analyzed by Northern blotting. Half-lives were determined as previously described (32). Briefly, target signal was normalized against control for the corresponding time and plotted on a graph. Slopes of exponential regressions were used to determine the first order decay kinetics to determine mRNA half-lives. Band mobility was determined by plotting signal intensity against migration distance and measuring the mobility of the peak of the target mRNA relative to the peak of the internal control mRNA, with mobility of the target mRNA at the zero-hour time point set to 100 and mobility of the target mRNA after treatment with RNase H and oligo-dT to remove the poly(A)-tail set to 0.

## Acknowledgements

We thank members of the Lykke-Andersen lab, Tim Nicholson-Shaw, Cody Ocheltree, and Tiantai Ma for input on the manuscript. We thank Georg Stoecklin (Heidelberg University) for sharing the anti-p-Ser178-TTP antibody. This work was supported by National Institutes of Health (NIH) fellowship F32 GM128320-03 to A.C. and National Institutes of Health (NIH) grant R35 GM118069 to J. L.-A.

## Legends to Supplementary Figures

**Supplementary Figure S1.**
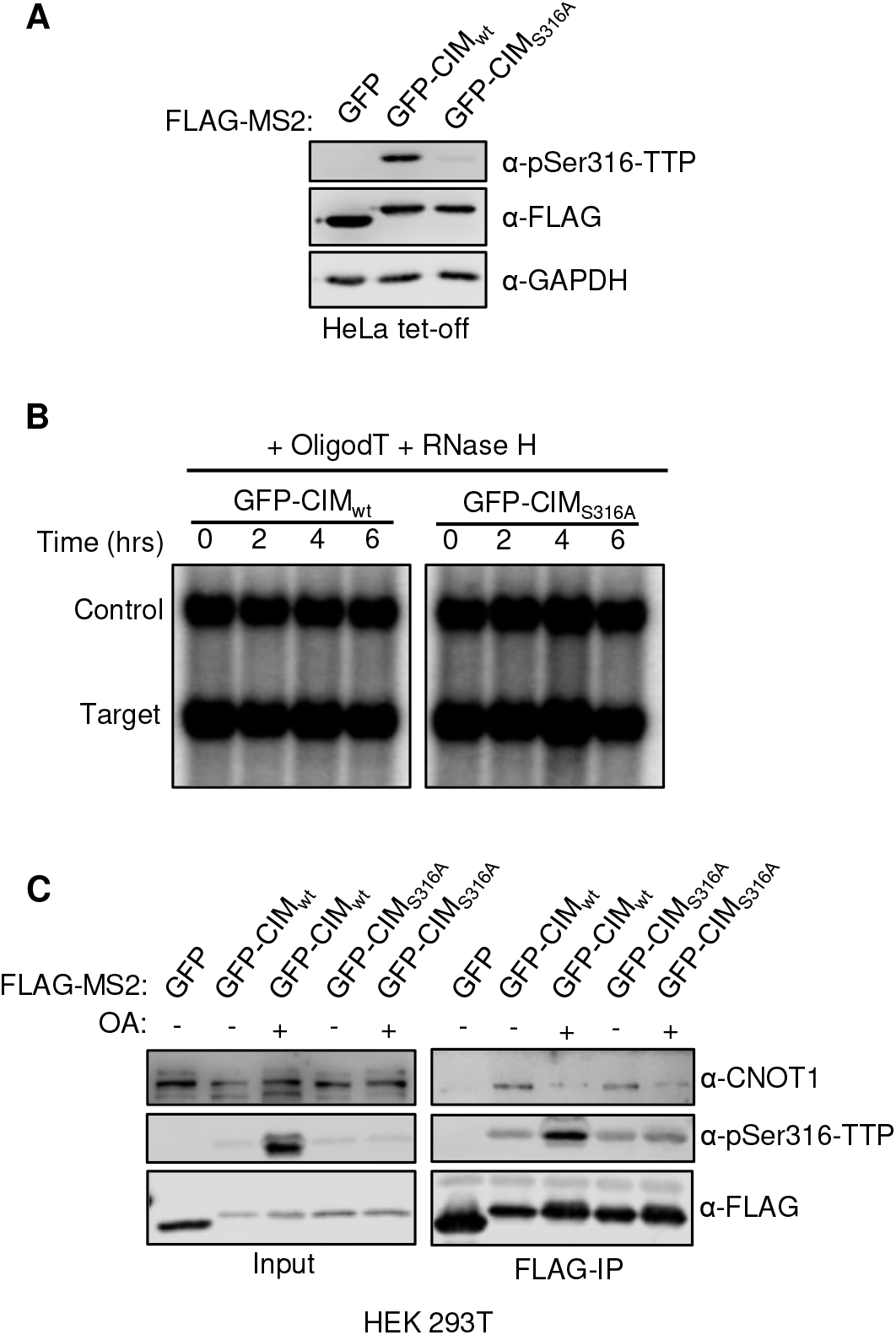
Effects of tethered TTP-CIM motif. (A) Western blots monitoring expression levels and phosphorylation of indicated FLAG-MS2-GFP fusion proteins in HeLa Tet-off cells. (B) Northern blot monitoring β-globin mRNA tethered to indicated MS2-GFP fusion proteins in HeLa tet-off cells and treated with oligo-dT and RNase H. (C) Western blots showing proteins co-immunoprecipitating (IP, right panels) with indicated FLAG-MS2-GFP fusion proteins from HEK293T cells treated with or without okadaic acid. Input samples corresponding to 2.5% of IPs are shown on the left.

**Supplementary Figure S2.**
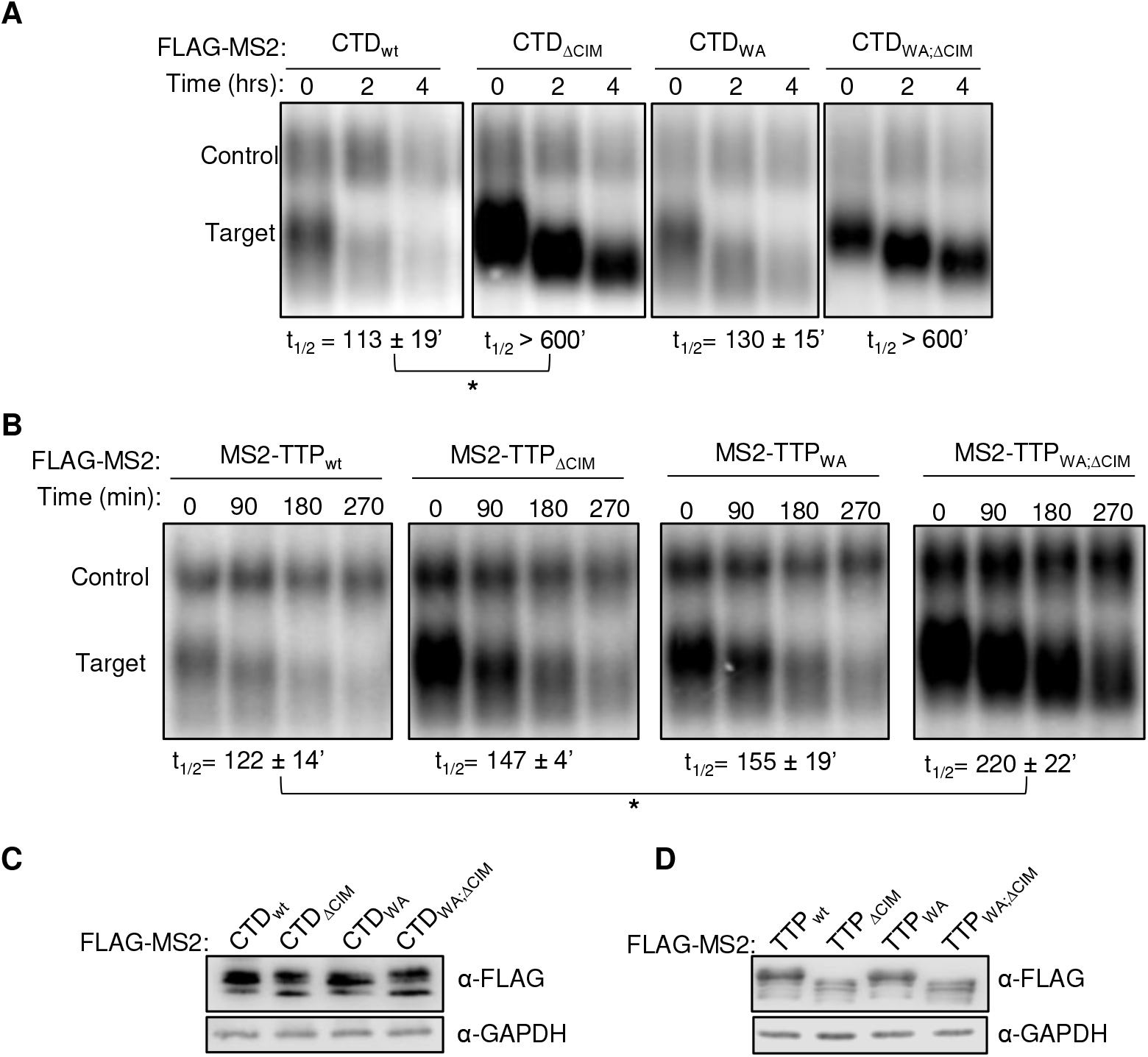
Cooperative activation of degradation by the TTP-CIM and conserved tryptophans. (A) Representative Northern blots monitoring the degradation of β-globin mRNA (Target) tethered to the indicated FLAG-MS2-CTD fusion proteins in HeLa Tet-off cells, graphed in Figure 2A. *: p<0.05; student’s two-tailed t-test. (B) Same as panel *A*, but monitoring degradation by MS2-TTP proteins graphed in Figure 2B. (C, D) Western blots monitoring expression levels of fusion proteins in panels *A* and *B*, respectively.

**Supplementary Figure S3.**
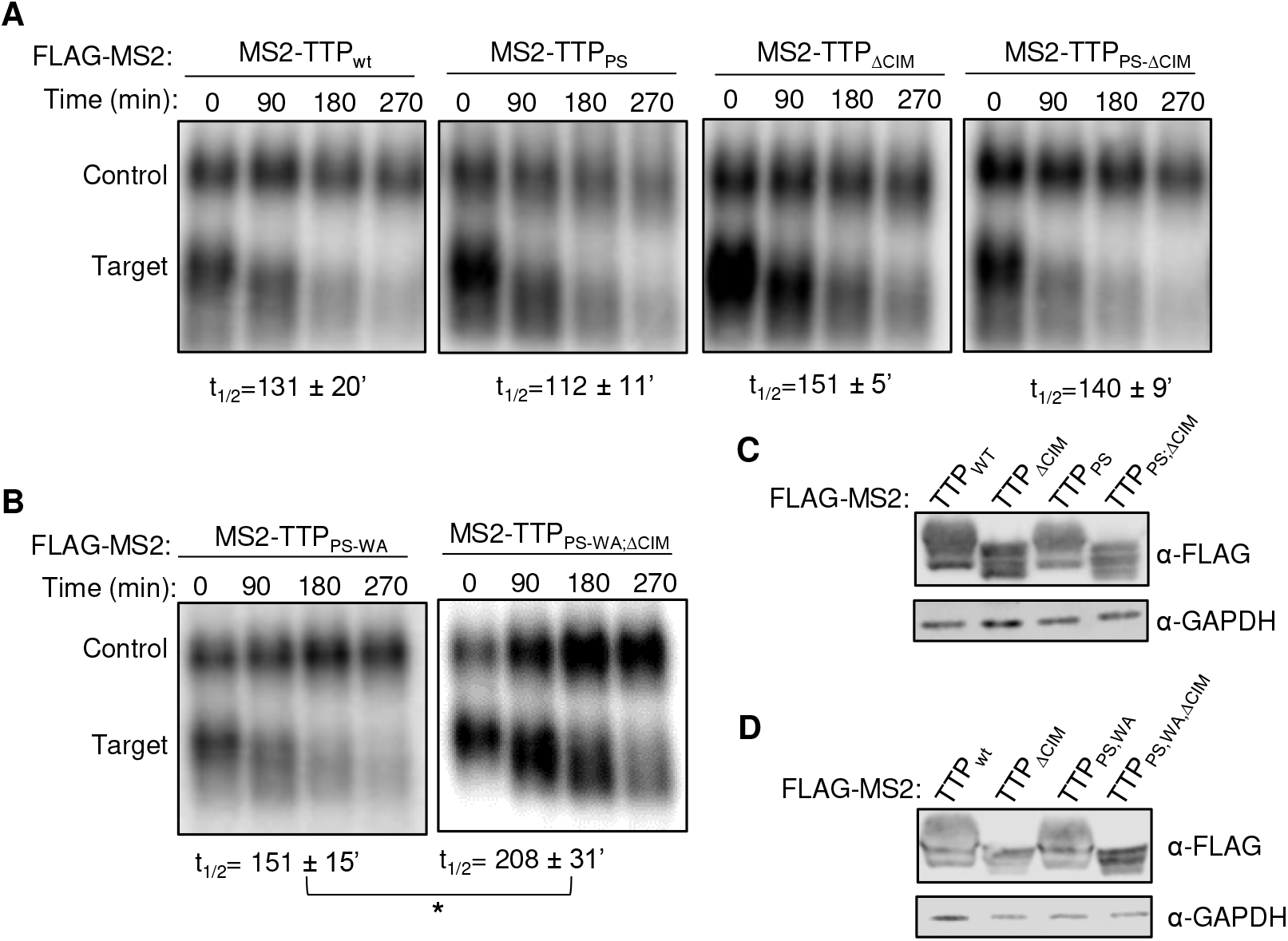
TTP tetraproline motifs are not rate-limiting for TTP-mediated mRNA decay. (A, B) Representative Northern blots monitoring the degradation of β-globin mRNA tethered to indicated MS2-TTP fusion proteins in HeLa Tet-off cells, graphed in Figures 3A and 3B, respectively. *: p<0.05; student’s two-tailed t-test. (C, D) Western blots monitoring MS2-TTP fusion protein expression levels in panels *A* and *B*, respectively.

**Supplementary Figure S4.**
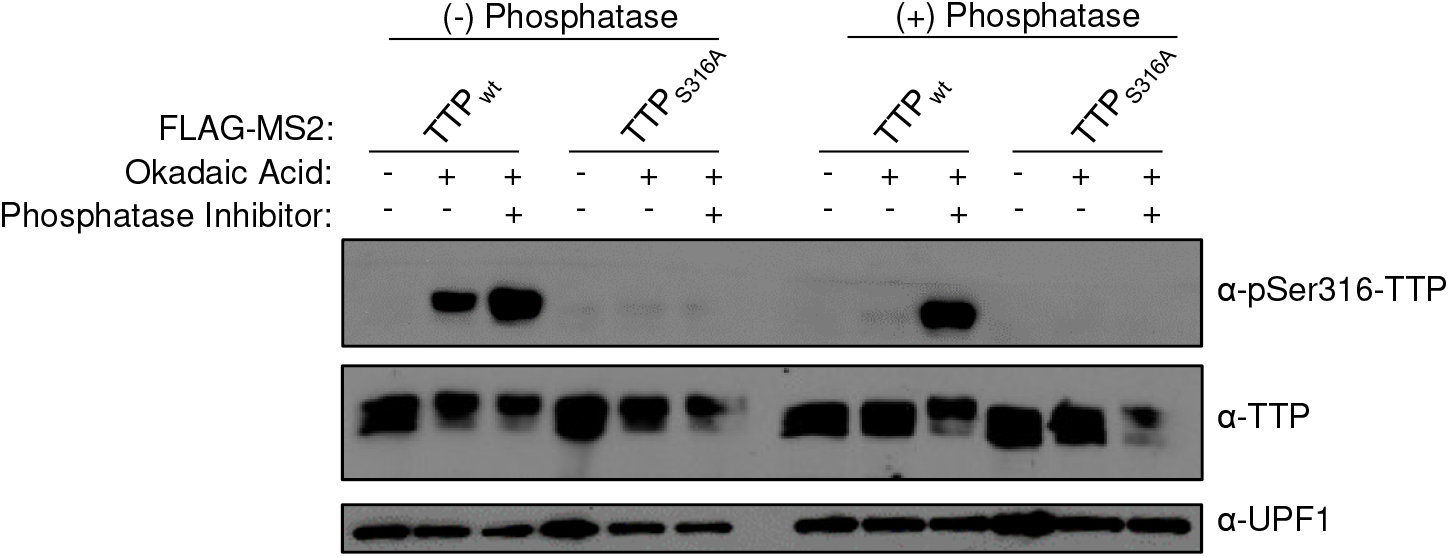
Validation of the anti-p-S316 TTP antibody. Western blot of indicated FLAG-MS2-TTP fusion proteins expressed in HEK293T cells incubated with or without okadaic acid (OA). Following cell lysis, samples were treated without (-, left panels) or with (+, right panels) calf-alkaline phosphatase in the presence or absence of phosphatase inhibitors (PhosphataseArrest I), prior to Western blotting using the anti-p-S316-TTP antibody as compared with TTP and UPF1 controls.

**Supplementary Figure S5.**
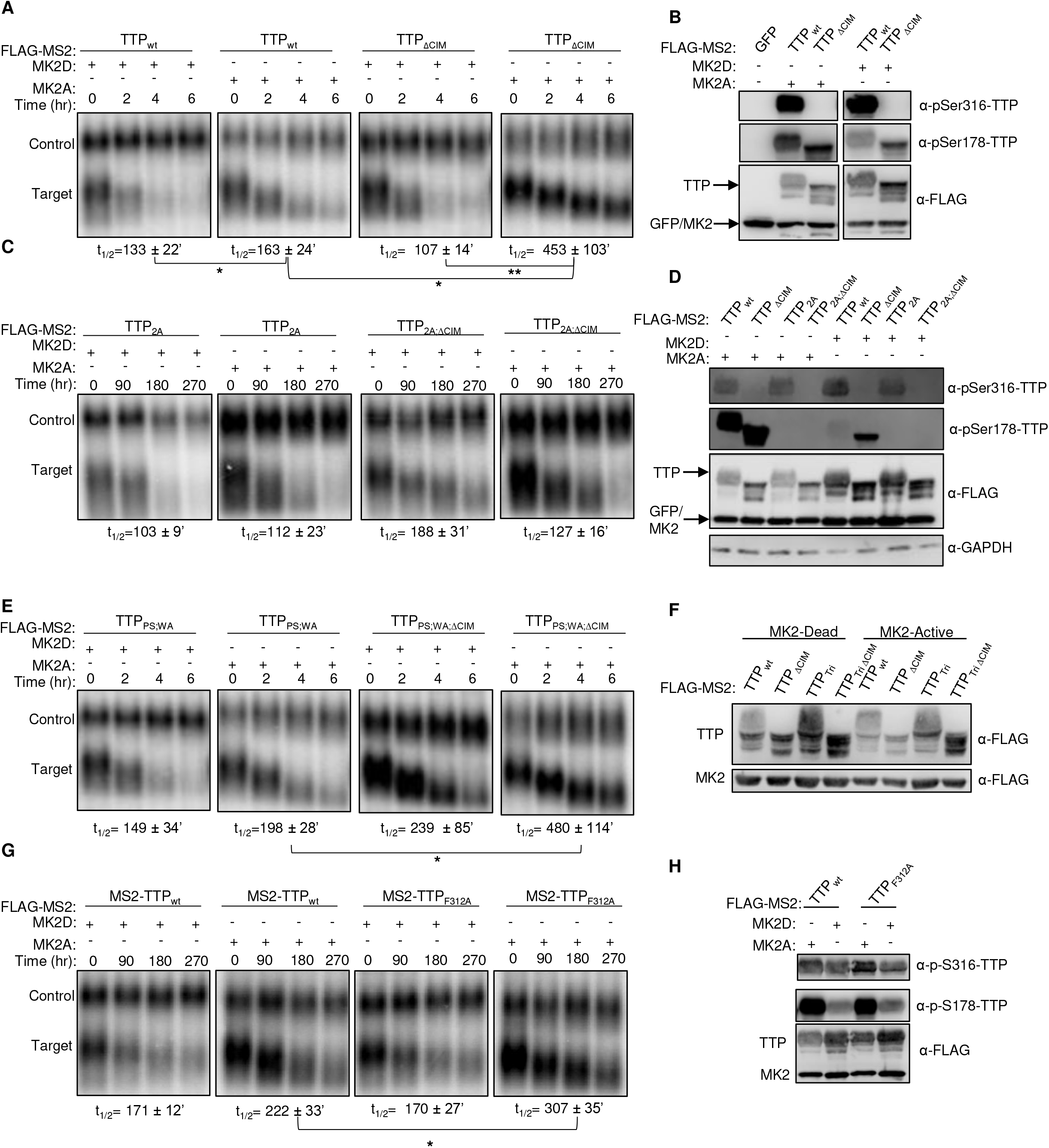
The TTP-CIM remains active in the presence of MK2. (A,C,E,G) Representative Northern blots monitoring degradation of β-globin mRNA tethered to indicated MS2-TTP fusion proteins in the presence of constitutive active or catalytically dead MK2 kinase in HeLa tet-off cells, some of which are graphed in Figure 5. (B,D,F,H) Western blots monitoring FLAG-MS2-TTP fusion protein phosphorylation and levels in the experiments in panels *A*, *C*, *E* and *G*.

**Supplementary Figure S6.**
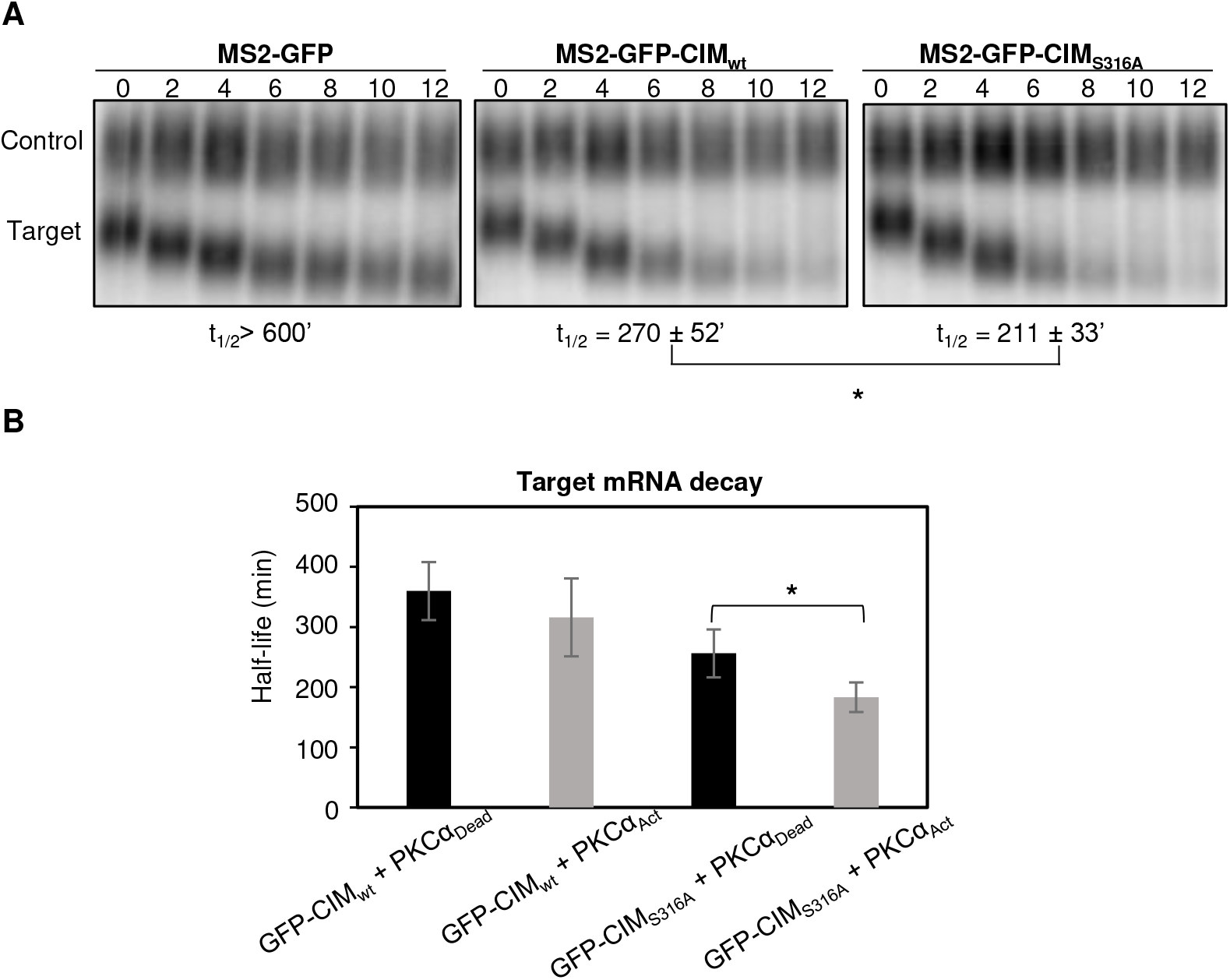
TTP-S316 phosphorylation inhibits mRNA deadenylation and decay. (A) Representative Northern blots monitoring degradation of β-globin mRNA tethered to indicated MS2-GFP fusion proteins, graphed in Figures 7A and 7B. (B) Bar graph showing calculated half-lives for β-globin mRNA tethered to MS2-GFP-CIM or MS2-GFP-CIM-S316A in the presence of co-expressed catalytic inactive (Dead) or constitutive active (Act) PKCα. *: p<0.05 (Student’s two-tailed t-test) (n=4).

---

**Supplementary Table 1.** List of FLAG-MS2-TTP mutant proteins

| FLAG-MS2: | TTP mutation(s): |
| --- | --- |
| CTD<sub>WA</sub> | Δ1-166; W255A,P257A |
| CTD<sub>WA;ΔCIM</sub> | Δ1-166; W255A,P257A; Δ304-319 |
| TTP<sub>WA</sub> | W31A,W61A,W255A,P257A |
| TTP<sub>PS</sub> | P64-66S; P191-193S |
| TTP<sub>WA;ΔCIM</sub> | W31A,W61A,W255A,P257A; Δ304-319 |
| TTP<sub>PS;ΔCIM</sub> | P64-66S; P191-193S; Δ304-319 |
| TTP<sub>PS;WA;ΔCIM</sub> | P64-66S; P191-193S; W31A,W61A,W255A,P257A; Δ304-319 |
| TTP<sub>S316A</sub> | S316A |
| TTP<sub>2A</sub> | S52A,S178A |

